# Prevention and treatment of noise-induced hearing loss and cochlear synapse degeneration by potassium channel blockers *in vivo*

**DOI:** 10.1101/2024.06.04.597382

**Authors:** Hong-Bo Zhao, Li-Man Liu, Ling Mei, Auraemil T Quinonez, Rafael A Roberts, Xiaoling Lu

## Abstract

Noise can induce hearing loss. In particularly, noise can induce cochlear synapse degeneration leading to hidden hearing loss, which is the most common type of hearing disorders in the clinic. Currently, there is no pharmacological treatment, particularly, no post-exposure (i.e., therapeutic) treatment available in the clinic. Here, we report that systematic administration of K^+^ channel blockers before or after noise exposure could significantly attenuate NIHL and synapse degeneration. After systematic administration of a general K-channel blocker tetraethylammonium (TEA), the elevation of auditory brainstem response (ABR) thresholds after noise-exposure significantly reduced, and the active cochlear mechanics significantly improved. The therapeutic effect was further improved as the post-exposure administration time extending to 3 days. BK channel is a predominant K^+^ channel in the inner hair cells. Systematic administration of a BK channel blocker GAL-021 after noise exposure also ameliorated hearing loss and improved hearing behavioral responses tested by acoustic startle response (ASR). Finally, both TEA and GAL-021 significantly attenuated noise-induced ribbon synapse degeneration. These data demonstrate that K^+^-channel blockers can prevent and treat NIHL and cochlear synapse degeneration. Our finding may aid in developing therapeutic strategies for post-exposure treatment of NIHL and synapse degeneration.

**Significance Statement:** Noise is a common deafness factor affecting more 100 million people in the United States. So far, there is no pharmacological treatment available. We show here that administration of K^+^ channel blockers after noise exposure could attenuate noise-induced hearing loss and synapse degeneration, and improved behavioral responses. This is the first time to real the K^+^ channel blockers that could treat noise-induced hearing loss and cochlear synaptopathy after noise exposure.

## Introduction

Noise can cause hearing loss and damages in the auditory system. High-intense noise can cause irreversible degeneration of spiral ganglion neurons (SGN) and hair cells leading to permanent threshold shift (PTS) in hearing. Noise also can induce temporal threshold shift (TTS) without apparent hair cell loss but having extensive loss of synaptic connections (ribbon synapses) between inner hair cells (IHCs) and auditory nerves (ANs) with a delayed degeneration of SGNs and their central projections and eventually lead to hidden hearing loss (HHL) (Kujawa and Liberman, 2009, 2015; Lin et al., 2011; Kobel et al., 2017; Boero et al., 2018). It is estimated that more than 100 million people have noise-induced hearing loss, which accounts for ∼10% of patients in the otolaryngology clinics (Hind et al., 2011; Kumar et al., 2012). However, except physical protections such as using earplugs, there are no pharmacological drugs available currently for treatment of noise-induced hearing loss (NIHL) and cochlear synaptopathy.

In previous studies, many drugs and antagonists and agonists of different cell signaling pathways, such as neurotrophins, TrkB agonists, and glucocorticoids, were tested for prophylactic and therapeutic interventions against NIHL (Han et al., 2015; Harrop-Jones et al., 2016; Poduslo and Curran, 1996; Ingersoll et al., 2020; Fernandez et al., 2021). Load noise also can increase the generation of reactive oxygen species (ROS). However, although therapeutics that target the free-radical pathways have shown some promises (He et al., 2021), none of these compounds is currently approved against NIHL by FDA. Moreover, most of previous studies mainly focused on noise-induced hair cell and SGN loss; only a few recent studies investigated noise-induced cochlear synapse degeneration (Yu et al., 2018; Hu et al., 2020; Ingersoll et al., 2020; Rouse et al., 2020; Seist et al., 2020; Fernandez et al., 2021). In addition, most of previous studies administrated drugs in pre-exposure (preventive) manners; only a few studies did post-exposure administration to test therapeutic effect against NIHL or cochlear synaptopathy (Shen et al., 2007; Bao et al., 2013; Yu et al., 2018; Dhukhwa et al., 2019; Ingersoll et al., 2020; Seist et al., 2020; Fernandez et al., 2021). Finally, the drug development and intervention have been challenged by the delivering method. Some drugs have been shown to have effects in the local administration (e.g., via the posterior semicircular canal) but not in the systematic administration (Poduslo and Curran, 1996; Fernandez et al., 2021), snice the blockade of crossing the blood-labyrinth barrier (BLB).

Noise can stimulate hair cells and neurons over-activating to increase K^+^ efflux elevating extracellular K^+^ concentration leading to K^+^-excitotoxicity, which can cause hair cell and cochlear synapse degeneration. Our previous study demonstrated that application of high-extracellular K^+^ in the *in vitro* preparation could cause cochlear ribbon synapse degeneration as shown in noise-induce HHL and that application of K^+^ channel blockers could attenuate this K^+^-excitotoxicity induced ribbon synapse degeneration (Zhao et al., 2021). In this study, we report that systematic administration of K^+^ channel blockers *in vivo* before or after noise exposure significantly attenuated NIHL and cochlear synapse degeneration. This study opens a new avenue for developing efficacious treatments to NIHL and cochlear synaptopathy. In particularly, the post-exposure treatment currently is unavailable but urgently required in the clinic.

## Materials and Methods

### Animal preparations

Adult male CBA/CaJ mice (8-16 weeks old, Stock No: 000654, The Jackson Lab) were used. Mice were housed in a quiet individual room in basement with regular 12/12 light/dark cycle. The background noise level in the mouse room at mouse hearing range (4-70 kHz) is <30 dB SPL. All procedures and experiments followed in the use of animals were approved by the University of Kentucky’s Institutional Animal Care & Use Committee (UK: 00902M2005) and Yale University’s Institutional Animal Care & Use Committee (Yale: 2022-20463) and conformed to the standards of the NIH Guidelines for the Care and Use of Laboratory Animals.

### Noise exposure and drug administrations

Mice were awake in a small cage under loud-speakers in a sound-proof chamber and exposed to white-noise (95-98 dB SPL) for 2 h, one time. The sound pressure level and spectrum in the cage were measured prior to placement of the animal (Zhao et al., 2021, 2022; Liu et al., 2023). For each time of noise exposure, 2-4 mice were exposed and randomly separated to treatment groups and control group to ensure age-matched control and background noise control among the experimental groups.

For pre-exposure treatment group, mice were intraperitoneally injected with drugs before 2 h of noise exposure. For post-exposure treatment group, mice were intraperitoneally injected immediately after noise exposure or one time per day for 3 days in the post-exposure 3-day groups. For general K^+^ channel blocker tetraethylammonium (TEA) (Cat#86614, Sigma-Aldrich, USA), the intraperitoneal injection (35 mg/kg) was administrated before or after noise exposure. For a specific BK channel blocker GAL-021 (Cat#22609, Cayman Chemical, Ann Arbor, MI, USA), only the post-exposure intraperitoneal injection (10 mg/kg) was performed, one time per day for 3 days. For the noise control group, mice were injected with the same amount of saline vehicle after noise exposure.

### Auditory brainstem response and distortion product otoacoustic emission recordings

ABR was recorded by use of a Tucker-Davis R3 workstation with ES-1 high frequency speaker (Tucker-Davis Tech. Alachua, FL) (Zhu et al., 2013, 2015; Mei et al., 2017; Zhao et al., 2022; Liu et al., 2023). ABR was recorded by stimulation with clicks in alternative polarity and tone bursts (4-40 kHz) from 80 to 10 dB SPL in a 5 dB step in a double-walled sound isolation room. The signal was amplified (50,000x), filtered (300-3,000 Hz), and averaged by 500 times. The ABR threshold was determined by the lowest level at which an ABR can be recognized. ABR was recorded at 1-2 day before noise exposure and post-exposure day 1, 3, 7, 14, and 28 to assess the threshold shift.

Distortion product otoacoustic emission (DPOAE) was recorded by two pure tones (f_1_ and f_2_, f_2_/f_1_ =1.22), which were simultaneously delivered into the ear. The test frequencies were presented by a geometric mean of f_1_ and f_2_ [f_0_ = (f_1_ x f_2_)^1/2^]. The intensity of f_1_ (I_1_) was set at 5 dB SPL higher than that of f_2_ (I_2_). The responses were averaged by 150 times (Zhu et al., 2013, 2015; Zhao et al., 2022; Liu et al., 2023).

### Acoustic startle response recording

Acoustic startle response (ASR) was recorded by use of a Startle II System (Kinder Scientific Inc, San Diego, CA). ASR was evoked by a series of white-noise pulses (25-ms duration) from 80 to 110 dB SPL in a 10 dB step.

### Immunofluorescent staining and confocal microscopy

Both of the treated cochlea and the control cochlea in the same mouse were stained in parallel to minimize variations of the performance. As described in our previous reports (Zhao and Yu, 2006; Liu and Zhao, 2008; Zhao et al., 2021, 2022; Liu et al., 2023), the mouse cochlea was freshly isolated, and the temporal bone was removed. The otic capsule was opened. The round window membrane was broken by a needle and a small hole was also made at the apical tip of the cochlea. Then, the cochlea was incubated with 4% paraformaldehyde in PBS for 0.5-1 h. After fixation, the cochlea was dissected in the normal extracellular solution (NES) (142 NaCl, 5.37 KCl, 1.47 MgCl_2_, 2 CaCl_2_, 10 HEPES in mM, pH 7.4). The basilar membrane with the organ of Corti (auditory sensory epithelium) was collected for staining.

The isolated cochlear sensory epithelia were incubated in a blocking solution (10% goat serum and 1% BSA of PBS) with 0.1% Triton X-100 for 30 min. Then, the epithelia were incubated with primary antibodies for CtBP2 (1:500, mouse anti-CtBP2 IgG1, #612044, BD Bioscience) and GluR (1:500, mouse anti-GluR2 IgG2a, #MAB397, Millipore Corp.) in the blocking solution at 4°C overnight. After washout, the epithelia were incubated with corresponding second Alexa Fluor antibody (1:400, gout anti-mouse IgG1 Alexa Fluor 568, A-21124 and goat anti-mouse IgG2a Alexa Fluor 488, A-21131, Thermo Fisher Sci) at room temperature (23 °C) for 2 hr. Before washout, 4’, 6-diamidino-2-phenylindole (DAPI, 50 μg/ml, D1306; Thermo Fisher Sci) was added to visualize the cell nuclei. After incubation for 2 min, the sensory epithelia were washed by PBS for 3 times and whole-mounted on the glass-slide (Zhao et al., 2021).

The stained epithelia were observed under a Nikon A1R confocal microscope system. CtBP2 labeling and DAPI staining were observed under a 561 nm laser with 570-620 nm emission filter and a 405 nm laser with a 425-475 nm emission filter, respectively. In each cochlea, at least 5 fields were scanned along the cochlear sensory epithelium from the apical to the basal. In each field, serial sections were taken along the Z-axis from the bottom to apical surface of the epithelium with a 0.25 μm step. All fluorescence microscopy was imaged using a Nikon 100x (N/A: 1.45) Plan Apro oil objective at 0.25 μm/pixel resolution (Zhao et al., 2021).

### Quantitative analysis and image presentation

As descripted in our previous publications (Zhao et al., 2021), we first generated the maximum intensity projection of z-stack images for each field of view. Then, the numbers of positive labeling for CtBP2 (ribbons) and GluR in the IHC region and the OHC region were separately accounted by NIS Elements AR Analysis software (Nikon). The numbers of ribbons and GluRs per IHC in each field of view was calculated and then averaged among the observed fields in each cochlea.

### Statistics and Reproducibility

Each experiment was repeated at least 4 times to verify the reproducibility. Data were plotted by SigmaPlot (Systat Software) and statistically analyzed by Excel (Microsoft Office 360) or SPSS (SPSS Inc., Chicago, IL). Error bars represent SEM. Parametric and nonparametric data comparisons were performed using one-way ANOVA or Student t tests after assessment of normality and variance. The threshold for significance was α = 0.05. Bonferroni *post hoc* test was used in ANOVA.

## Results

### Noise-induced hearing loss and attenuation by systematic application of a K^+^ channel blocker TEA before and after noise exposure

Fig. 1 shows noise-induced hearing loss that could be attenuated by systematic administration of a K^+^-channel blocker tetraethylammonium (TEA) before and after noise exposure. Mouse ABR thresholds after noise exposure were significantly increased (Fig. 1). Before noise exposure, ABR thresholds for click, 8, 16, 24, 32, and 40 kHz tone stimulations were 30.5±0.8, 27.7±1.04, 16.6±1.34, 26.6±1.26, 36.6±1.43, and 37.9±1.68 dB SPL, respectively, in the noise group, 33.4±0.75, 29.1±0.82, 15.9±0.94, 26.3±0.97, 35.6±0.9, and 37.8±1.71 dB SPL, respectively, in the pre-exposure TEA administration group (Pre-TEA group), and 33.1±0.59, 30.6±0.55, 15.8±0.49, 27.1±0.85, 35.8±0.89, and 36.7±1.03 dB SPL, respectively, in the post-exposure TEA administration group (Post-TEA group). There were no significant differences in ABR thresholds among these groups (Fig. 1E). After noise exposure, ABR thresholds for click, 8, 16, 24, 32, and 40 kHz tone stimulations at post-exposure day 1 (P1) in the noise group were increased to 91.6±3.23, 77.5±2.65, 76.8±1.17, 81.6±1.49, 84.5±1.83, and 90.2±1.59 dB SPL, respectively (Fig. 1F). However, ABR thresholds for click, 8, 16, 24, 32, and 40 kHz tone stimulations at P1 were 75.3±1.85, 69.4±3.32, 70.3±1.25, 81.6±1.09, 82.5±2.19, and 87.8±1.51 dB SPL, respectively, in the pre-TEA group and 70.0±2.76, 65.8±2.91, 68.8±1.57, 78.5±1.61, 86.0±1.90, and 88.1±1.75 dB SPL, respectively, in the post-TEA group (Fig. 1F). In comparison with those in the noise group, ABR thresholds at click, 8 kHz, and 16 kHz in both pre- and after-TEA groups were significantly reduced by 10-20 dB SPL (P< 0.01 or 0.05, one-way ANOVA with a Bonferroni correction) (Fig. 1F). At P28 (Fig. 1G), ABR thresholds for click, 8, 16, 24, 32, and 40 kHz tone stimulations were 52.2±1.77, 47.2±1.82, 45.0±2.54, 58.4±1.49, 65.3±1.91, and 72.8±2.14 dB SPL, respectively, in the pre-TEA group and 52.3±1.99, 50.4±1.99, 51.7±2.25, 64.6±1.90, 71.9±1.16, and 78.1±1.34 dB SPL, respectively, in the post-TEA group. In comparison with those in the noise group, which were 63.6±1.87, 57.0±1.76, 63.4±1.16, 71.4±0.88, 76.6±1.20, and 79.3±1.06 dB SPL for click, 8, 16, 24, 32, and 40 kHz tone stimulations, respectively (Fig. 1G), the ABR thresholds in both pre- and post-TEA groups were significantly reduced by 5-20 dB SPL (P< 0.01 or 0.05, one-way ANOVA with a Bonferroni correction) (Fig. 1G).

**Fig. 1.**
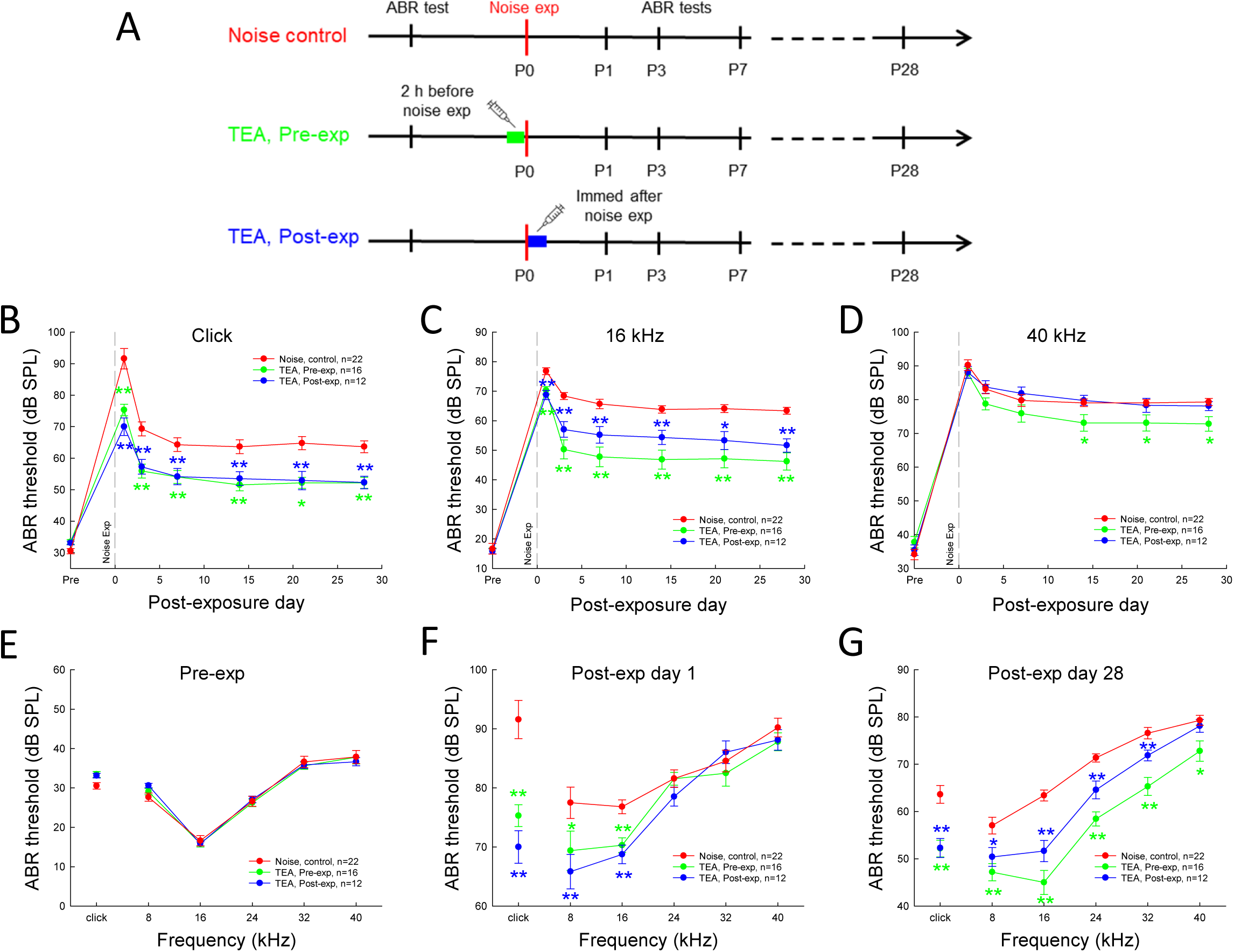
Rescuing of hearing by systematic administration of TEA before and after noise exposure. **A:** Schematical drawings of experimental procedures for ABR recording, noise exposure, and systematic administration of TEA before and after noise exposure. Mice at 3 months old were exposed to 98 dB SPL white noise for 2 h. Before noise exposure, ABR is recorded to define mouse pre-exposure hearing. A K^+^ channel blocker TEA (35mg/kg) was intraperitoneally injected to mice at 2 h before noise exposure (pre-TEA group), or immediately after exposure for one time (post-TEA group). The noise group was injected with the same saline vehicle after noise exposure. The day of noise exposure was set as post-exposure day 0 (P0). ABR was recorded at P1, P3, P7, P14, P21, and P28. **B-D:** Dynamic changes of ABR thresholds after noise exposure in the noise group, the pre-TEA group, and the post-TEA group. ABR thresholds in both pre- and post-TEA groups were significantly improved in comparison with those in the noise group. **E-G:** ABR thresholds of the noise group, the pre-TEA group, and the post-TEA group at pre-exposure, P1, and P28. Asterisks represent the statistical significance referring to the noise group. *: P <0.05, **: P<0.01, one-way ANOVA with a Bonferroni correction.

### Improvement of superior-threshold hearing function by systematic administration of TEA

Systematic administration of TEA before and after noise exposure also improved superior-threshold hearing. Fig. 2 shows that the amplitudes of ABR evoked by 90 dB SPL click stimulation were significantly increased in both pre- and post-TEA groups in comparison with those in the noise group (Fig. 2B-H). The peak of wave I at P28 in both pre- and post-TEA groups were almost 2 times larger than that the noise group (Fig. 2B-C). The negative peak of wave VI also had large increase in the pre- and post-TEA groups (Fig. 2B&H).

**Fig. 2.**
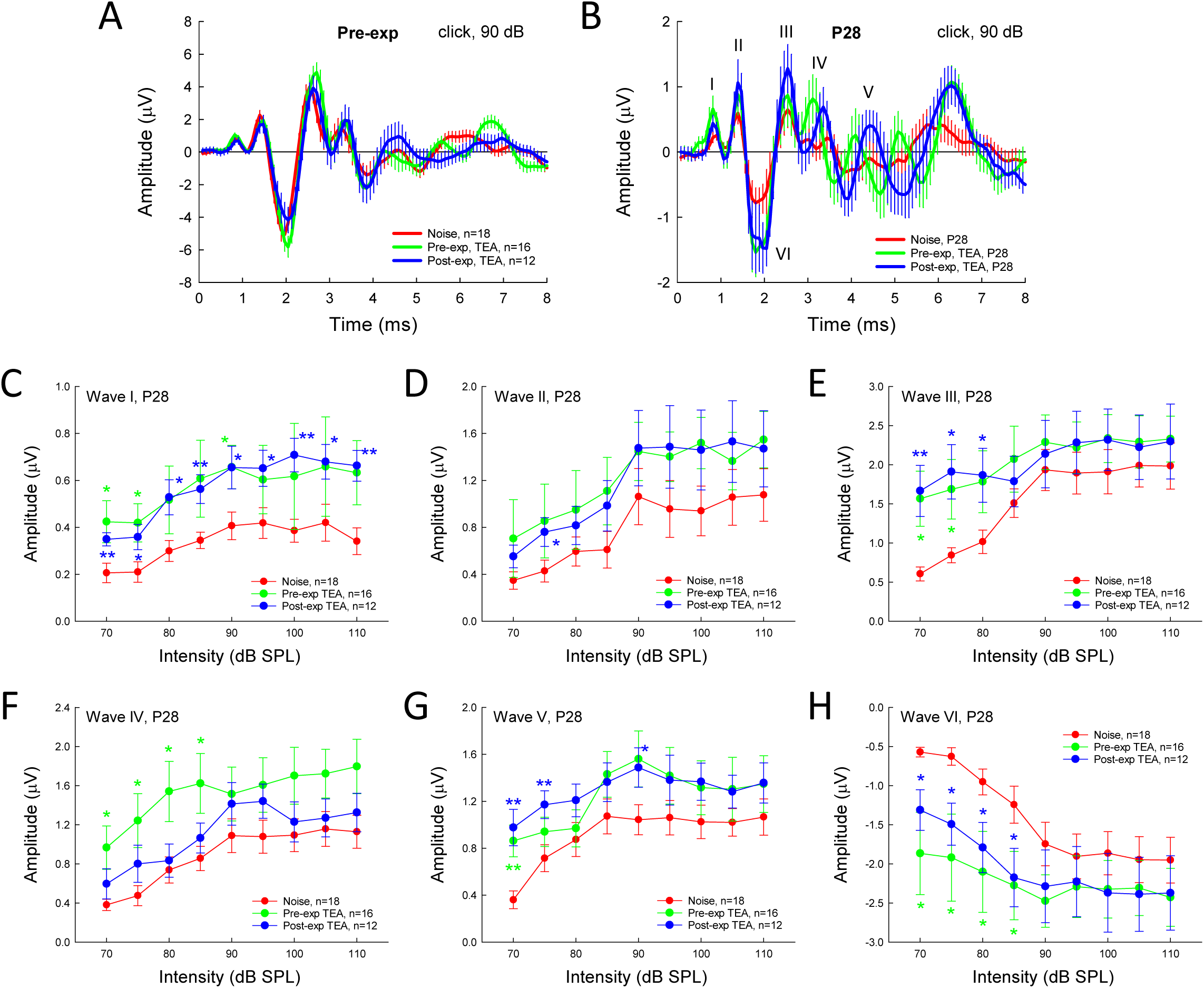
The effects of pre- and post-exposure administrations of TEA on amplitudes of ABR. The ABR was evoked by 90 dB SPL click stimuli. **A-B**: The averaged waveforms of ABRs at pre-exposure and P28 after noise exposure in the noise group, and the pre- and post-TEA groups. **C-H**: The quantitative measurements of ABR amplitudes in the noise, pre- and post-TEA groups at P28. Asterisks represent the statistical significance referring to the noise group. *: P <0.05, **: P<0.01, one-way ANOVA with a Bonferroni correction.

### Improvement of active cochlear amplification by systematic administration of TEA

Systematic administration of TEA before and after noise exposure also attenuated the noise-induced impairment in the active cochlear amplification (Fig. 3). Before noise exposure, there was no significantly difference in DEOAEs among the noise group, pre- and post-TEA groups (Fig. 3D). After noise-exposure, DPOAEs in the noise group significantly reduced (Fig. 3A&E-F). In comparison with the pre-exposure level, the reduction could be up to 40 dB at the high intensity (Fig. 3F). Pre- or post-exposure administration of TEA significantly attenuated this reduction (Fig. 3B-C&E). Referring to that in the noise group, the DPOAEs in both pre- and post-TEA groups were improved by 10-15 dB (P< 0.01 or 0.05, one-way ANOVA with a Bonferroni correction) (Fig. 3E-F).

**Fig. 3.**
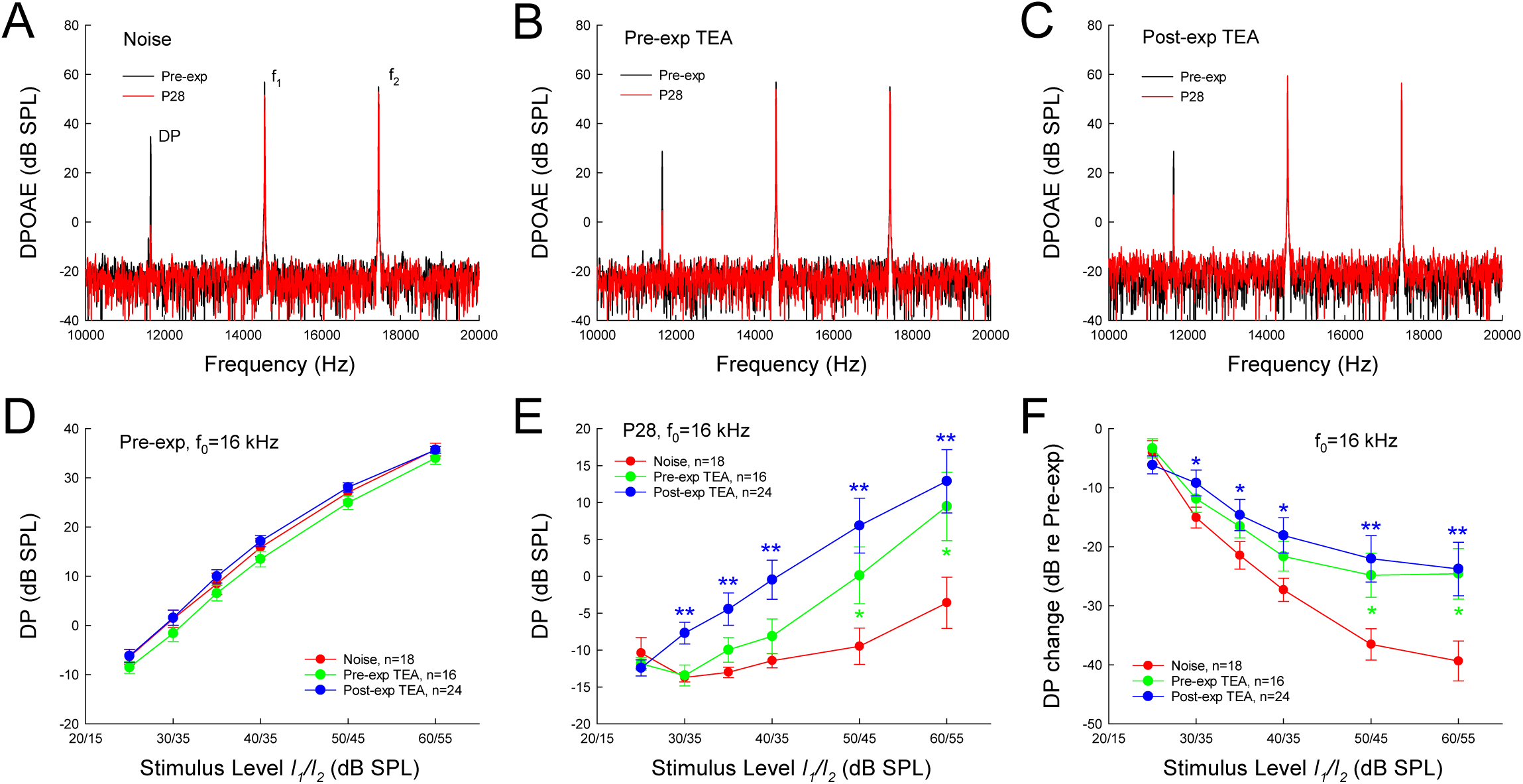
Reduction of DPOAE after noise exposure and rescue by pre- and post-exposure administration of TEA. **A-C**: The spectra of acoustic emission in the noise, pre-TEA, and post-TEA groups at pre-exposure and P28 after noise exposure. Acoustic emission was evoked by two pure tones: *f_0_* = (*f_1_* x *f_2_*)^1/2^ =16 kHz, *f_2_/f_1_* =1.22, *I_1_/I_2_* = 60/55 dB SPL. **D-F**: Changes of DPOAEs at pre-exposure and P28 in the noise, pre-TEA and post-TEA groups. Panel **F** represents DP changes referring to the DP level at pre-noise exposure in each group. Asterisks represent the statistical significance referring to the noise group. *: P <0.05, **: P<0.01, one-way ANOVA with a Bonferroni correction.

### Improvement of the therapeutic efficacy by prolonging administration of TEA

We further tested whether prolonging administration time of TEA could increase therapeutic efficacy. Fig. 4 shows that after post-exposure administration of TEA for 3 days (the post-TEAx3d group), ABR thresholds at P28 had a significant recovery (Fig. 4B&C). Comparing with those in the noise group without TEA treatment, ABR thresholds in the post-TEAx3d group at P28 had ∼20 dB improvement in all tested frequency range (P< 0.01, one-way ANOVA with a Bonferroni correction) (Fig. 4C). In comparison with ABR thresholds in the single-doze post-TEA group (or post-TEAx1d group), improvement of the ABR thresholds in the post-TEAx3d group was also significant (P< 0.01 or 0.05, one-way ANOVA with a Bonferroni correction) (Fig. 4D&E). The improvement (referring to the noise group) was increased from 20-40% in the post-TEAx1d group to 50-70% in the post-TEAx3d group, in particularly, in the high-frequency range (Fig. 4D). Fig. 4E further shows the fold changes of ABR thresholds in the post-TEAx3d group referring to the post-TEAx1d group. The improvement of therapeutic efficacy increased as frequency increased. The therapeutic effects could be improved by ∼10-fold at high frequencies (Fig. 4E).

### Attenuation of noise-induced hearing loss by BK channel blocker

BK channel is a dominant K channel in the cochlea and mainly express in the inner hair cells (Fig. 5D, Pyott and Duncan, 2016). We further tested whether the BK channel blocker could attenuate NIHL. A small molecule GAL-021 is a specific BK channel blocker and capable of penetrating through the brain-blood barrier (BBB) (McLeod et al., 2014; Roozekrans et al., 2014, 2015; Golder et al., 2015). Fig. 5 shows that systematic administration of GAL-021 after noise exposure for 3 days significantly ameliorated NIHL. At day 28 after noise exposure, ABR thresholds in the GAL-treatment group (GALx3d) were 53.0±2.13, 47.0±1.86, 50.5±1.89, 53.0±3.27. 60.0±2.89, and 72.5±1.86 dB SPL for click, 8, 16, 24, 32, and 40 kHz tone stimulations, respectively (Fig. 5C). In comparison with those in the noise group, the ABR thresholds in the GALx3d group were significantly attenuated by 15-25 dB SPL (P< 0.01 one-way ANOVA with a Bonferroni correction) (Fig. 5C). If referring to the increase in ABR thresholds in the noise group, the recovery ratio of ABR thresholds in the GALx3d group was ∼50%, as the same as the recovery ratio in the post-TEAx3d group (Fig. 5E).

**Fig. 4.**
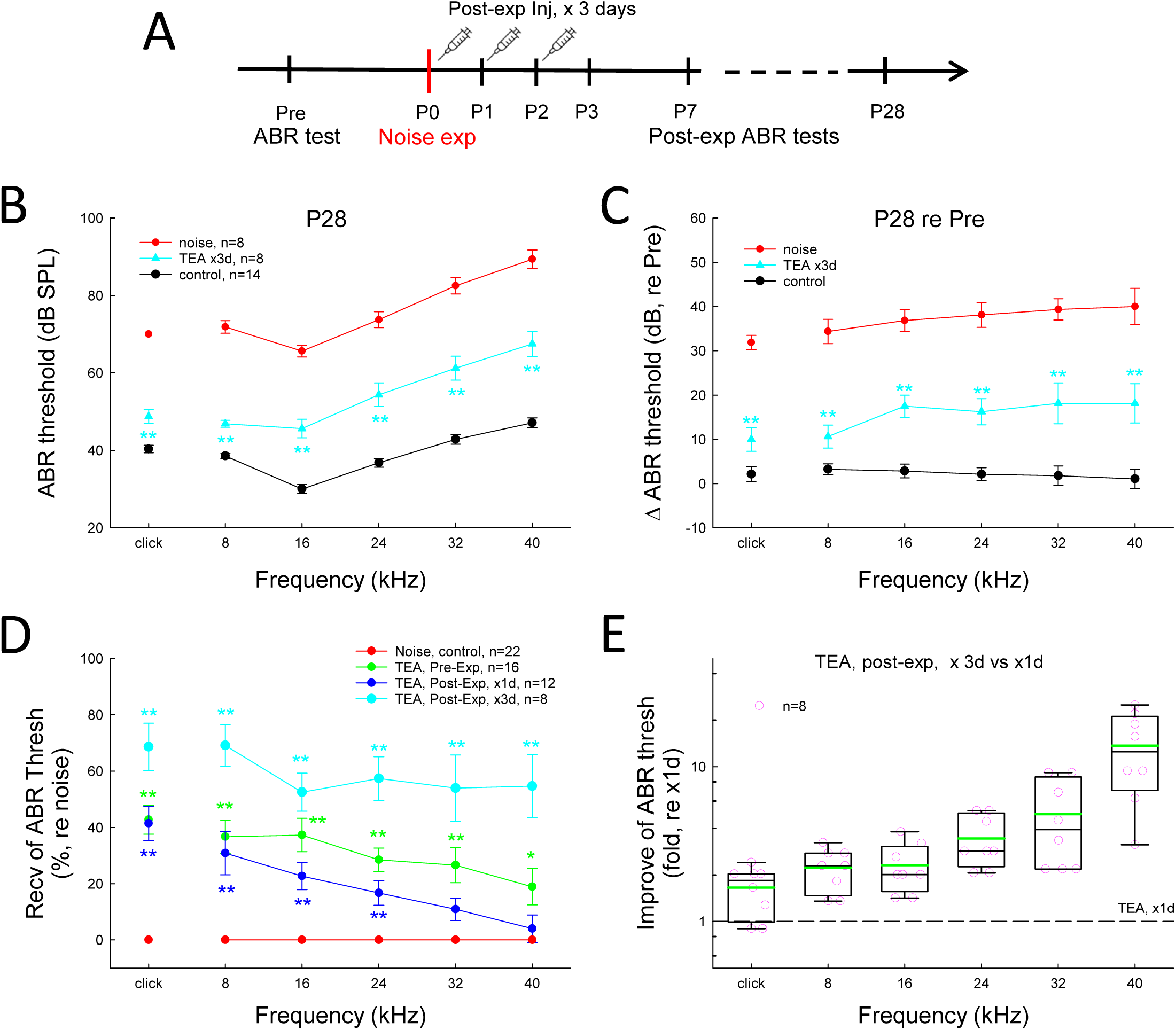
Improvement of therapeutic effect by systematic administration of TEA for 3 days after noise exposure. **A**: the schematical drawing of experimental designing and procedure. Mice were intraperitoneally injected by TEA after noise exposure, one time per day for 3 days (post-TEAx3d group). **B-C**: ABR thresholds and changes at P28 in the noise group, post-TEAx3d group, and control group. To be visible clearly, only comparisons between the noise and post-TEAx3d groups are marked. **D**: Recovery of ABR thresholds of pre- and post-exposure administration of TEA referring to those in the noise group. The recovery of ABR thresholds is normalized to those in the noise group at P28. **E**: The improvement of the ABR thresholds in the post-TEAx3d group referring to those for post-exposure administration of TEA for 1 day, i.e., post-TEA or post-TEAx1d group (indicated by a dash-line). The ABR thresholds in the post-TEAx3d group have apparent improvement in comparison with those in the post-TEAx1d group. *: P <0.05, **: P<0.01, one-way ANOVA with a Bonferroni correction (panel **B-D**), or 2-tail t test (panel **E**).

**Fig. 5.**
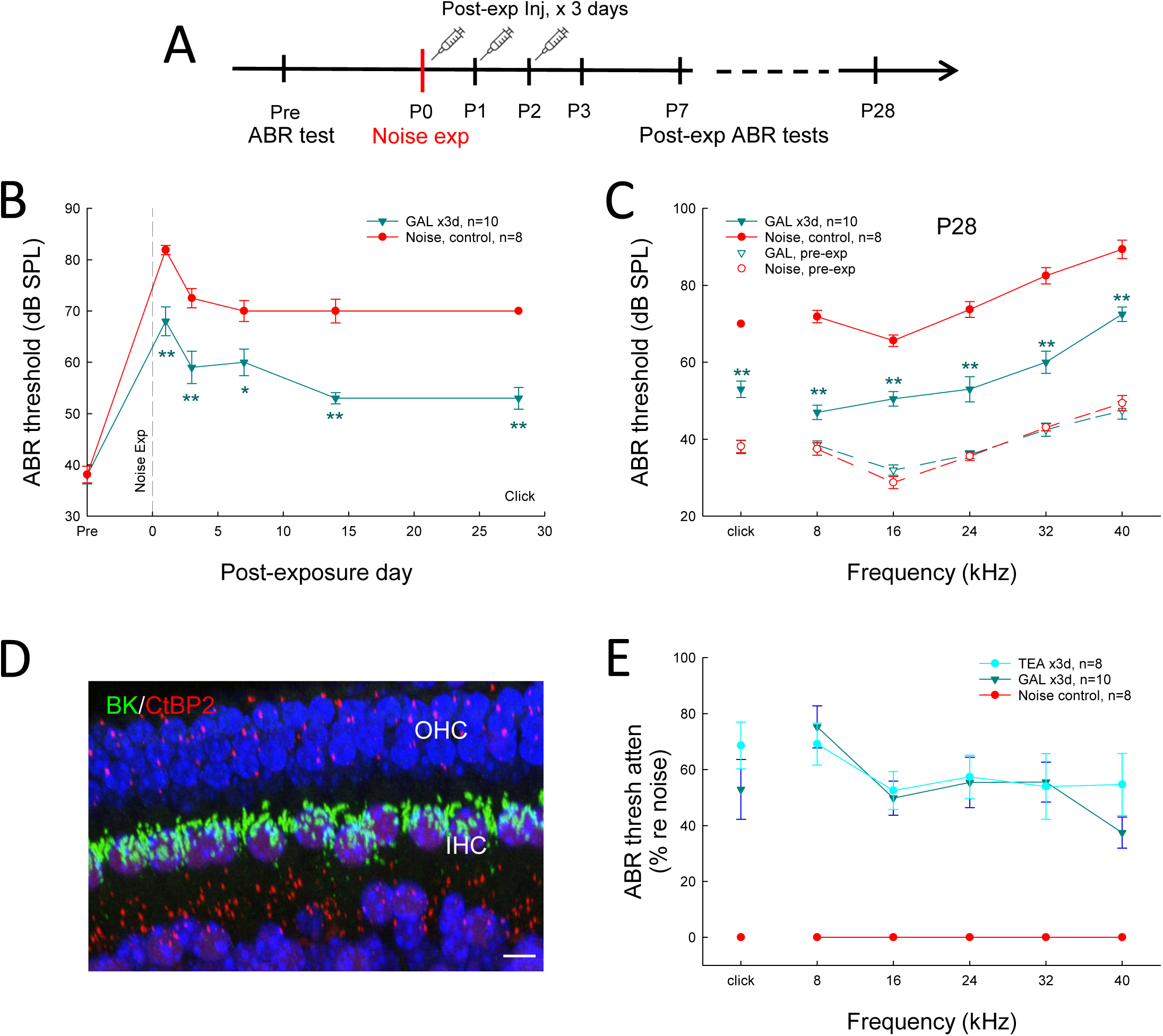
Attenuation of noise-induced hearing loss by systematic administration of a BK channel blocker GAL-021 after noise exposure. GAL-021 is a specific BK channel blocker and were intraperitoneally injected into mice (10 mg/kg) after noise exposure by 1 time per day for 3 days (GALx3d group). **A**: A schematic drawing for experimental procedure. **B**: Dynamic changes of ABR thresholds after noise exposure with GAL-021 injections. **C**: ABR thresholds in the GALx3d group at P28 were significantly reduced in comparison with those in the noise group. Dashed lines represent the ABR thresholds of the noise group and GALx3d group before noise exposure. **D**: BK channel expression in the cochlea by immunofluorescent staining of the cochlear sensory epithelium with the whole-mountain preparation. IHC, inner hair cell; OHC, outer hair cell. Scale bar: 10 µm. **E**: Comparison of treating effects of GAL-021 and TEA. There is no significant difference in the recovery of ABR thresholds between the GALx3d group and the post-TEAx3d group. *: P<0.05, **: P<0.01, 2-tail t test in panel **B** and one-way ANOVA with a Bonferroni correction in panel **C**.

GAL-021 also attenuated the acute increase of ABR thresholds after noise exposure. Fig. 5B shows that the ABR thresholds of click in the GALx3d group at P1, P3, and P7 were significantly reduced by ∼10 dB SPL in comparison with those in the noise group (P< 0.01, t test).

### Improvement of hearing function measured by acoustic startle response

We further assessed the therapeutic effect by behavioral testing. Acoustic startle responses (ASRs) in the noise, GALx3d, and normal control groups were recorded. Fig. 6 shows that noise exposure significantly reduced ASR; the ASR in the noise group at P40-45 was significantly smaller than that in the control group. ASRs in the GALx3d group were significantly improved, like the ASR recorded in the control group (Fig. 6). There was no significant difference between the GALx3d group and the control group (Fig. 6C&D).

**Fig. 6.**
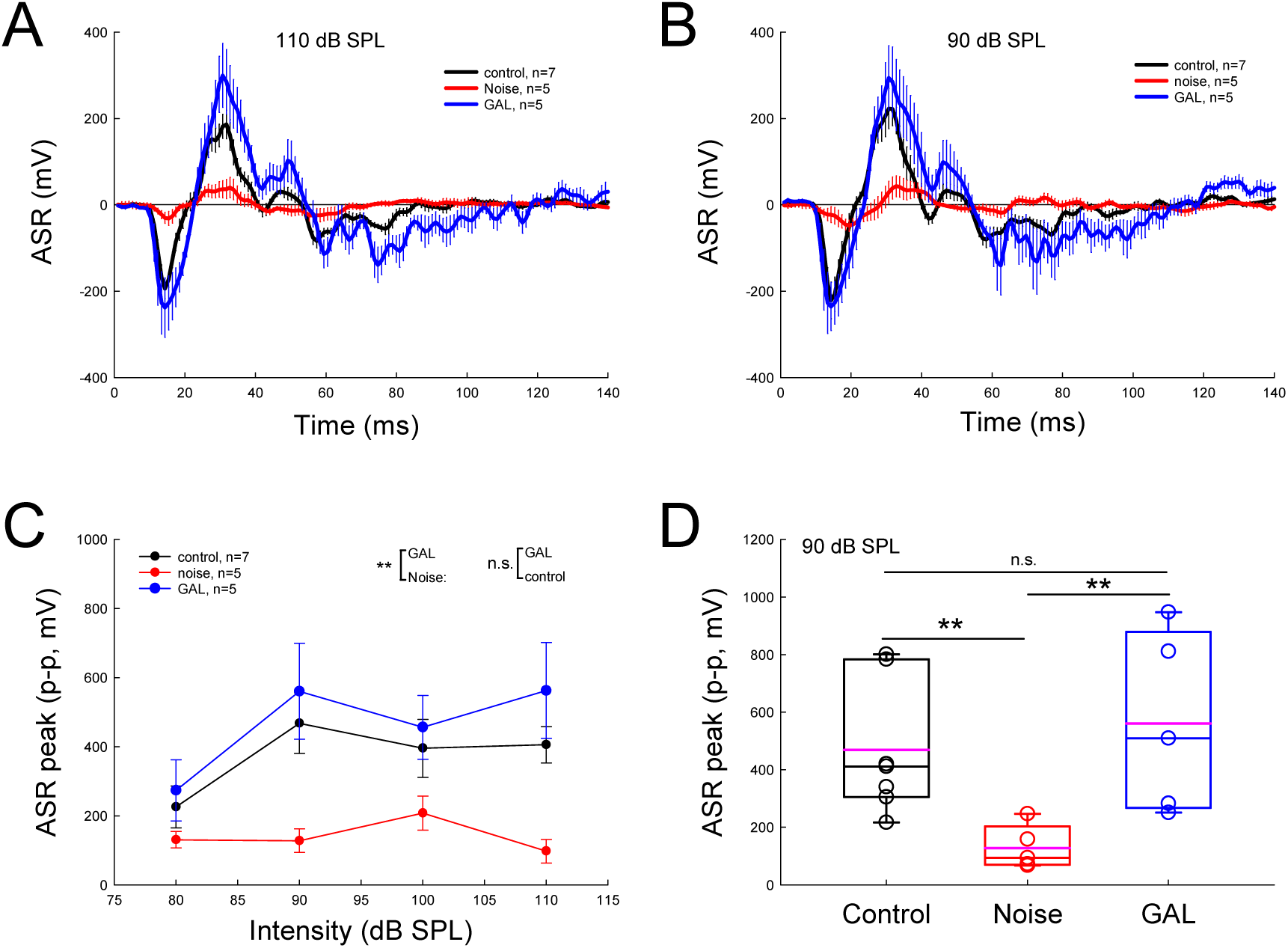
Recovery of hearing function measured by acoustic startle response (ASR) after systematic post-exposure administration of GAL-021 for 3 days. ASR was evoked by a series of white-noise pulses (25 ms width, 80-110 dB SPL, 10 dB step), and was measured at post-noise exposure day 40-45. The control group was not given to noise exposure. **A-B**: The averaged ASR waveforms recorded from the control group (black), noise group (red), and GAL-021 treated group (blue) evoked by 90 and 110 dB white-noise pulses. Error bars represent SE. **C**: The amplitudes of ASRs (peak to peak) measured from the control, noise, and GAL groups. There is significant difference between the noise group and control group (P< 0.01, F test). However, there is no significant difference between the GAL group and control group (P=0.22, F test). **D**: The peak of ASR evoked by 90 dB SPL white-noise pulse in the control, noise, and GAL groups. There is a significant reduction of ASR in the noise group (**: P < 0.01, one-way ANOVA with a Bonferroni correction). However, there is no significant difference in ASRs between the control group and the GAL group (P=0.59, one-way ANOVA).

### Attenuation of IHC ribbon synapse degeneration by TEA

We further examined the ribbon synapse degeneration after noise exposure and attenuation by the administration of TEA (Fig. 7). After noise exposure, the numbers of ribbons, GluRs, and synapses at the apical, middle, and basal turns were significantly reduced in comparison with those in the control group (Fig. 7B-D). The degeneration appeared apparent at the middle and basal turns, in where the numbers of ribbons, GluRs, and synapses were reduced by ∼50% (Fig. 7B-D). However, after administration of TEA, the degenerations were significantly attenuated, especially at the apical and middle turns (Fig. 7B-D). For example, the numbers of synapses at the apical, middle, and basal turn were significantly increased from 5.77±0.46, 5.82±0.20, and 1.68±0.28 per IHC, respectively, in the noise group to 7.39±0.56, 8.05±0.40, and 2.79±0.33/IHC, respectively, in the pre-TEA group and 7.94±0.37, 7.01±0.23, and 2.72±0.23/IHC, respectively, in the post-TEA group (P< 0.01 or 0.05, one-way ANOVA with a Bonferroni correction) (Fig. 7D). The ratios of synaptic ribbons and GluRs referring to total ribbons and GluRs were also significantly increased after administrations of TEA (Fig. 7E&F). The ratio of synaptic ribbons at the apical, middle, and basal turns were significantly increased from 57.3±3.70, 68.5±2.40, and 38.0±7.80%, respectively, in the noise group to 70.3±3.53, 76.9±2.61, and 50.9±4.94%, respectively, in the pre-TEA group and 76.2±2.45, 76.0±1.91, and 55.5±3.85%, respectively, in the post-TEA group (P< 0.01 or 0.05, one-way ANOVA with a Bonferroni correction) (Fig. 7E). The increases were by 10-20%. The ratios of synaptic GluRs were also significantly increased, in particularly, at the basal turn (Fig. 7F). At the basal turn, the ratio of synaptic GluRs was 58.3±6.55% in the noise group and significantly increased to 83.5±3.13 and 91.4±1.71% in the pre-TEA group and post-TEA group, respectively (P< 0.01 or 0.05, one-way ANOVA with a Bonferroni correction) (Fig. 7F); the increases were by 25-30%.

**Fig. 7.**
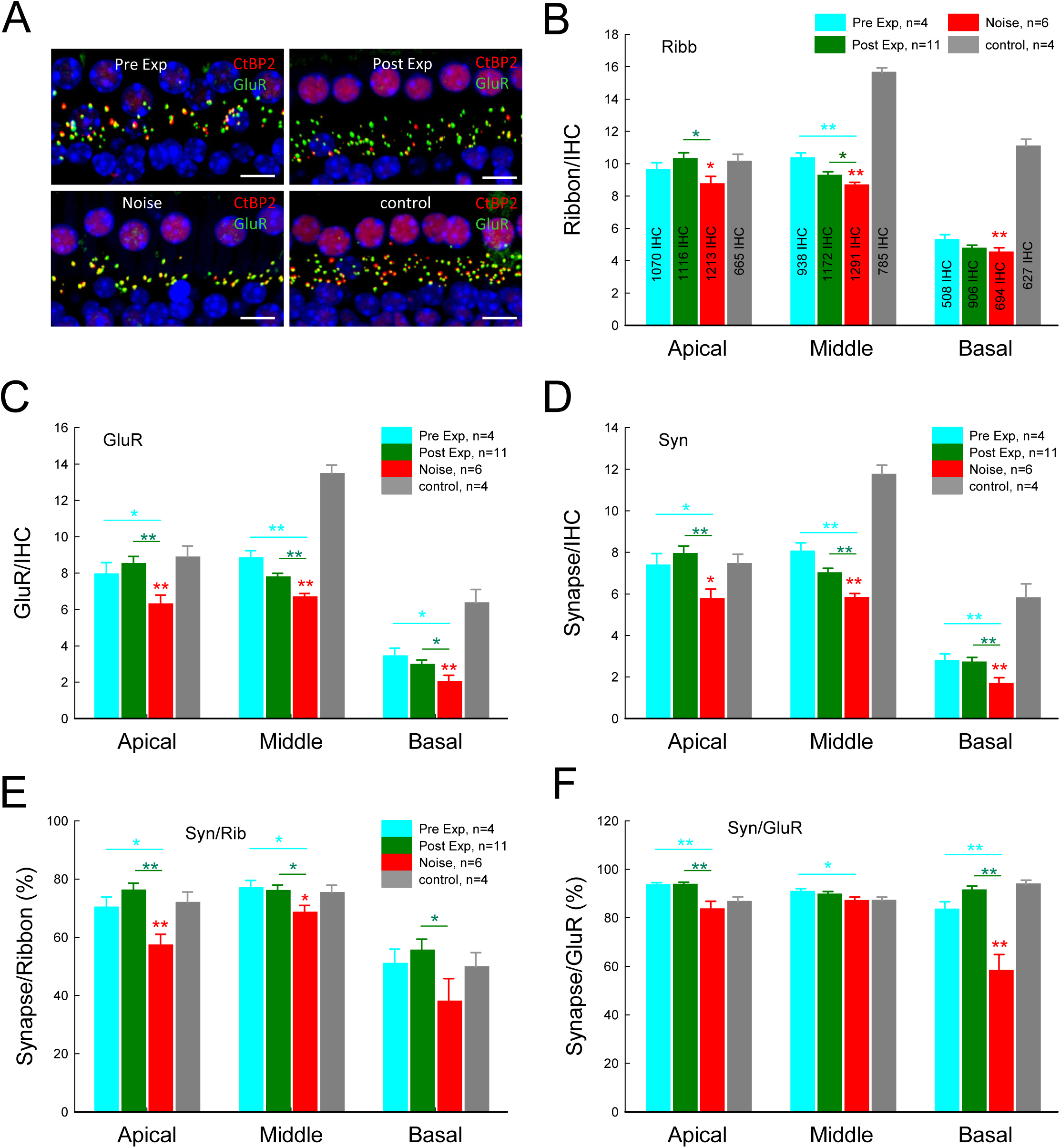
Noise exposure induced ribbon synapse degeneration and attenuated by TEA. **A**: Immunofluorescent staining for CtBP2 and GluR in the pre-TEA group, post-TEA (or post-TEAx1d) group, noise group, and control group. Mice were killed at post-exposure day 30-45. Scale bar: 10 µm. **B-D**: Noise exposure induced significant degeneration of ribbon synapses in comparison with the control group (indicated by red asterisks). However, the numbers of ribbons, GluRs, and synapses per inner hair cell (IHC) at the apical, middle, and basal turn in TEA injection groups are significantly increased comparing with those in the noise group. Numbers in the bars in panel **B** represent the total numbers of accounted IHCs in each group. **E-F**: Changes of ratios of synaptic ribbons and synaptic GluRs after noise exposure and TEA administrations. The ratio of synaptic ribbons or GluRs was calculated from normalized to the total numbers of ribbons or GluRs. Red asterisks represent significant changes between the noise group and control group. *: P<0.05, ** P < 0.01, one-way ANOVA with a Bonferroni correction.

Fig. 8 shows attenuation of ribbon synapse degeneration by post-exposure administration of GAL-021 for 3 days (GALx3d group). The attenuation appeared apparent at the apical and middle turns (Fig. 8B-D). In particularly, the numbers of ribbons, GluRs, and synapses at the apical turn in the GALx3d group were 10.9±0.37, 10.9±0.44, and 10.1±0.38/IHC, respectively, as the same as those in the control group, which were 10.4±0.48, 10.9±0.54, and 9.91±0.46/IHC, respectively (Fig. 8B-D). At the middle turn, the numbers of ribbons, GluRs, and synapses in the GALx3d group were 9.56±0.51, 9.34±0.45, and 8.71±0.44/IHC, respectively, and were still significantly higher than those in the noise group, which were 8.04±0.53, 7.72±0.46, and 6.89±0.45/IHC, respectively (P< 0.01 or 0.05, one-way ANOVA with a Bonferroni correction) (Fig. 8B-D). However, there was no significant attenuative effect of GAL-021 on ribbon synapse degeneration at the basal turn (Fig. 8B-D).

**Fig. 8.**
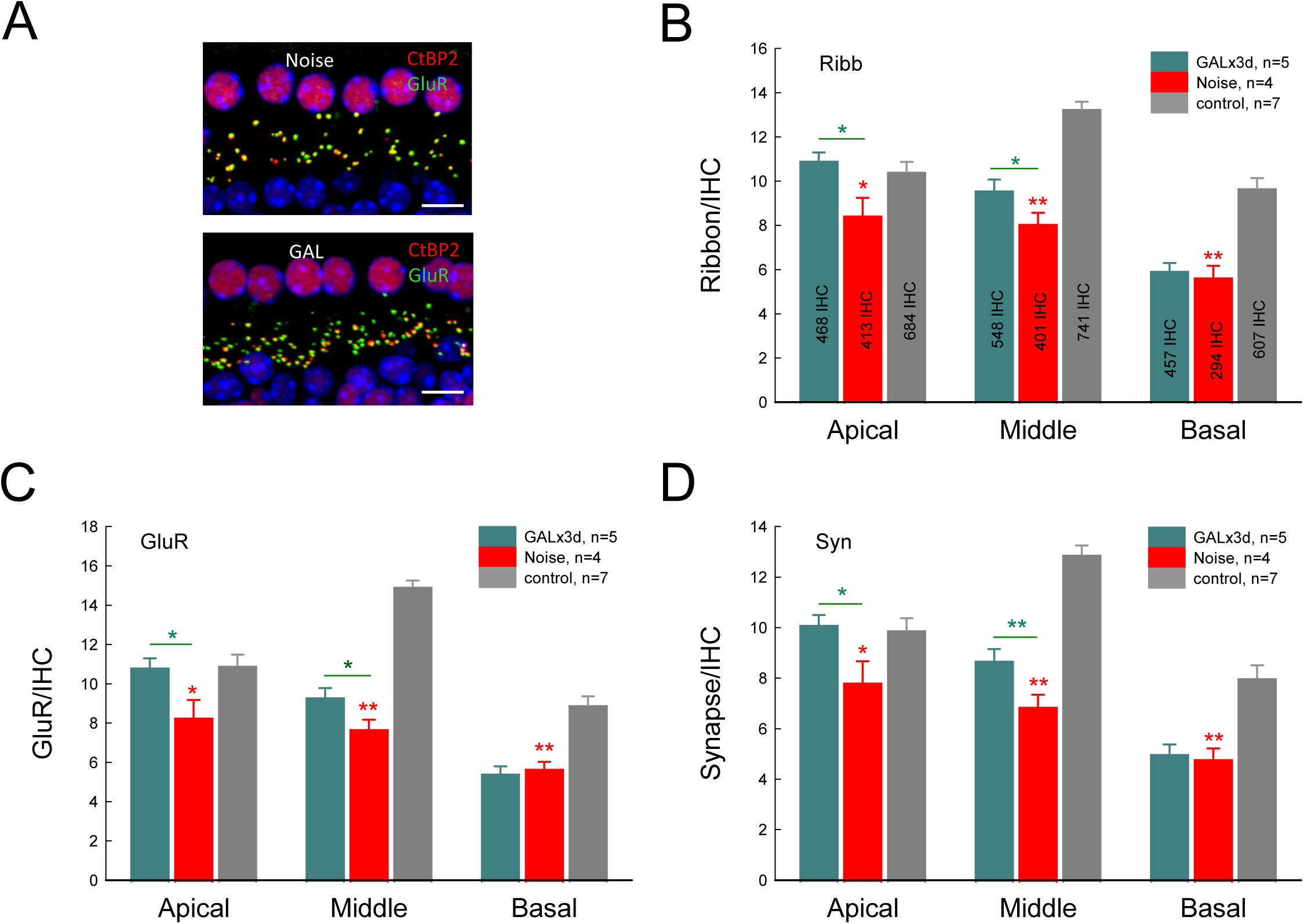
Attenuation to noise-induced IHC ribbon synapse degeneration by post-exposure administration of GAL-021 for 3 days. **A**: Immunofluorescent staining for CtBP2 and GluR in the GAL-021 administration and noise groups. Scale bar: 10 µm. **B-D**: Numbers of ribbons, GluRs, and synapses per IHC in the GALx3d group are significantly increased comparing with those in the noise group. Red asterisks indicate the significant reduction of ribbon synapses in the noise group in comparison with control group. Numbers in the bars in panel **B** represent the total numbers of accounted IHCs in each group. *: P<0.05, **: P<0.01, one-way ANOVA with a Bonferroni correction.

Like ABR thresholds improved as post-exposure administrative time extended to 3 days (Fig. 4), Fig. 9 shows that the numbers of ribbons, GluRs, and synapses per IHC in both post-TEAx3d and GALx3d groups were also significantly increased. Except ribbons at the apical and middle turns, GluRs and synapses in both post-TEAx3d and GALx3d groups were significantly increased at all apical, middle, and basal turns (Fig. 9A-C). In comparison with those in the TEAx1d group, the increases of synapse in the TEAx3d and GALx3d groups were by 120-200% (P< 0.01 or 0.05, one-way ANOVA with a Bonferroni correction) (Fig. 9D). The ratios of synaptic ribbons at the apical, middle, and basal turns also increased from 76.1±2.45, 76.0±1.91, and 55.5±3.85%, respectively, in the post-TEAx1d group to 91.9±1.36, 92.1±1.69, and 83.0±3.31%, respectively, in the post-TEAx3d group and 92.8±0.89, 91.6±1.10, and 84.0±2.13%, respectively, in the GALx3d group (P< 0.01 or 0.05, one-way ANOVA with a Bonferroni correction) (Fig. 9E). However, there were no significant changes in the ratio of synaptic GluRs among the post-TEAx1d, post-TEAx3d, and GALx3d groups, except the middle turn, at which the ratios of synaptic GluRs in both the post-TEAx3d and GALx3d groups were significantly increased from 89.7±1.18% in the TEAx1d group to 93.3±0.98% and 93.1±0.69%, respectively (P< 0.01 or 0.05, one-way ANOVA with a Bonferroni correction) (Fig. 9F). However, outer hair cells (OHCs) had no significant loss in the experimental groups (Fig. S1).

**Fig. 9.**
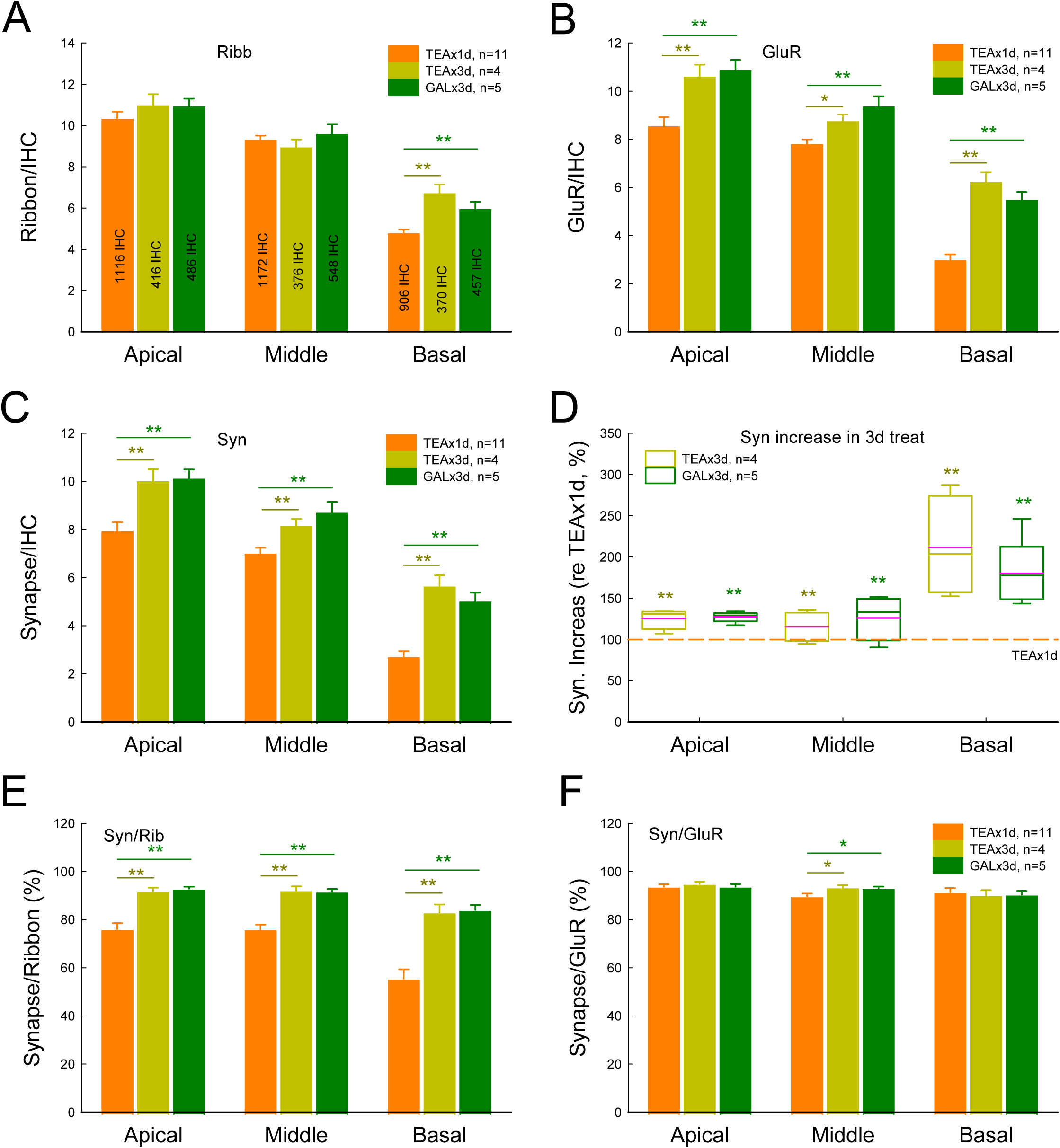
Improvement of attenuation of ribbon synapse degeneration by post-exposure administration of GAL-021 and TEA for 3days. **A-C**: Numbers of ribbons, GluRs, and synapses per IHC at the apical, middle, and basal turn for post-exposure administration of TEA for one day (post-TEAx1d) and 3 days (post-TEAx3d), and GAL-021 for 3 days (GALx3d). In comparison with those in the post-TEAx1d group, the numbers of ribbon, GluRs, and synapses per IHC at the middle and basal turns in both post-TEAx3d and GALx3d groups have significant improvements. Numbers in the bars in panel **A** represent the total numbers of accounted IHCs in each group. **D:** Improvement of synapses in the post-TEAx3d and GALx3d groups in comparison with those in the post-TEAx1d group. The synapse had large increase at the basal turn in the post-TEAx3d and GALx3d groups. **E-F**: Increments of synaptic ribbons and GruRs for 3 days treatment. The ratios of synapse/ribbon at the apical, middle, and basal turn in both post-TEAx3d and GALx3d groups were significantly increased in comparison with those in the post-TEAx1d group (penal **E**). However, synaptic GluRs only at the middle turn were significantly increased (penal **F**). *: P<0.05, **: P<0.01, one-way ANOVA with a Bonferroni correction.

## Discussion

In this study, we found that systematic administrations of K^+^ channel blockers before or after noise exposure significantly attenuated NIHL and cochlear synapse degeneration (Figs. 1, 2, and 7). The active cochlear amplification (Fig. 3) and superior threshold hearing function (Fig. 6) were also improved. The therapeutic effects became more apparent as the administration extended to post-exposure 3 days (Fig. 4, 8, and 9). The BK channel is a predominant K channel in the IHCs (Pyott and Duncan, 2016). A specific BK channel blocker GAL-021 also had a significant therapeutic effect on NIHL and attenuated ribbon synapse degeneration (Figs. 5, 6, 8, and 9). This *in vivo* study is consistent with our previous *in vitro* study (Zhao et al., 2021), demonstrating that K^+^ channel blockers could ameliorate NIHL and cochlear synaptopathy. This finding may aid in developing potential therapeutic strategies for post-exposure treatment of NIHL, which currently is not available but urgently required in the clinic.

The prophylactic and therapeutic effects of drugs on NIHL largely depend on delivering methods (i.e., local *vs* systematic), since the blockade of crossing the blood-labyrinth barrier (BLB). For example, although neurotrophins were extensively tested as therapeutic agents, the difficulty in targeting therapeutic levels of proteins to the brain or peripheral nerves has hampered their use (Poduslo and Curran, 1996). Also, it was reported that local administration (via the posterior semicircular canal) of TrkB agonists could regenerate synapses after noise-induced synaptopathy (Fernandez et al., 2021). However, the systematic administration only had an effect when given before noise exposure at high dose. It was reported that systematic administration of a small molecule bisphosphonate (zoledronate), which can penetrate the BLB, to mice after 24 h after noise exposure could partially regenerate IHC synapses (Seist et al., 2020). ABR threshold was not changed but the amplitude of its wave I was increased (Seist et al., 2020). Steroid dexamethasone was also reported to be partially effective in treating noise ototoxicity (Han et al., 2015; Harrop-Jones et al., 2016). However, randomized controlled trials reported contradictory outcomes (Crane et al., 2015). In this study, we found that systematic administration of K^+^ channel blocker TEA could attenuate NIHL (Figs. 1-4) and cochlear synaptopathy (Figs. 7-9). In particularly, we found that a specific BK channel blocker GAL-021, which can penetrate the brain-blood barrier (McLeod et al., 2014; Roozekrans et al., 2014, 2015; Golder et al., 2015), could significantly ameliorate NIHL (Figs. 5&6) and attenuate IHC ribbon synapse degeneration (Figs. 8&9). This is consistent with previous report that BK channel deficiency could increase resistance to NIHL (Pyott et al., 2007; Pyott and Duncan, 2016). Our finding further suggests that the BK channel has an important role in the prevention and protection from noise and that BK channels could be a potential target for prevention and treatment for NIHL and ribbon synapse degeneration.

The prophylactic and therapeutic effects of drugs on NIHL are also determined by administrative time, i.e., pre- *vs* post-noise exposure. Most of previous studies administrated drugs in pre-exposure or mixed (pre- plus post-exposure) manners; only a few studies did post-exposure administration to test therapeutic effect against NIHL or cochlear synaptopathy (Shen et al., 2007; Bao et al., 2013; Yu et al., 2018; Dhukhwa et al., 2019; Ingersoll et al., 2020; Seist et al., 2020; Fernandez et al., 2021). However, despite these statistics, pharmacological therapies for NIHL, particularly post-exposure treatments for HHL and synapse degeneration, currently are still not existent. In this study, we systematically administrated K^+^ channel blockers before or after noise exposure and found that both pre- and post-exposure systematic administrations of K^+^ channel blockers significantly attenuated NIHL and cochlear synaptopathy (Figs. 1-9). Our study may open a new avenue for developing efficient drugs for prevention and treatment of noise-induced synapse degeneration and deafness.

Our study also provides important information about mechanisms underlying NIHL and cochlear ribbon synapse degeneration, which currently is still unclear. It has been hypothesized that the cochlear synaptopathy may result from glutamate-excitotoxicity, i.e., acoustic overstimulation (noise exposure) induces excessive glutamate release leading to ribbon synapse excitotoxicity and degeneration (Liberman and Kujawa, 2017). However, previous experiments demonstrated that the presynaptic and postsynaptic components remained intact after perfusion of GluR agonists (Puel et al., 1991; Sebe et al., 2017). Moreover, the excess glutamate induced hair cell death, which was independent of damage of postsynaptic terminals (Sheets, 2017). Our previous study (Zhao et al., 2021) also demonstrated that application of GluR agonists could cause ribbon swelling but not degeneration and that application of GluR antagonist had no effect on ribbon degeneration. These results suggest that except for glutamate-excitotoxicity, other mechanisms also play an important role in NIHL and cochlear synaptopathy in HHL.

Noise exposure can elevate extracellular K^+^ concentration due to hair cell and neuron over-activation increasing K^+^ efflux. It is well-known that high extracellular K^+^ can produce excitotoxicity to cause hair cell and synapse degeneration. In our previous study (Zhao et al., 2021), we demonstrated that high extracellular K^+^ could cause IHC ribbon synapse degeneration as induced by noise exposure and applied K^+^ channel blockers could attenuate this ribbon synapse degeneration in the *in vitro* preparation (Zhao et al., 2021). In this *in vivo* study, we found that administration of K^+^-channel blockers could ameliorate NIHL (Figs. 1-5) and attenuate cochlear synapse degeneration (Figs. 7-9). This is consistent with our previous *in vitro* study, and further indicates that K^+^-excitotoxicity has an important role in the hair cell and synapse degeneration in NIHL.

It has been reported that age-related hearing loss at the early stage has apparent cochlear synapse degeneration pertaining to potassium channels in the cochlea and auditory pathway (Sergeyenko et al., 2013; Fernandez et al., 2015; Viana et al., 2015; Kujawa and Liberman, 2015; Parthasarathy and Kujawa, 2018; Peixoto et al., 2021). Our study revealed that K^+^-channel blockers could attenuate cochlear synapse degeneration (Figs. 7-9). Our study may also shed light on targeting age-related hearing loss and suggests that K^+^-channels may have a role in the age-related hearing loss. In addition, recently, it has been found that noise plays an important role in Alzheimer’s disease (AD) development and progressing (Cantuaria et al., 2021). This study may also open a new avenue for AD protection and prevention.

## Acknowledgement

This work was supported by NIH R01 DC 017025 and RF1 AG082216 to HBZ.

## Author Contributions

HBZ conceived the general framework of this study. LML, ML, ATQ, XL, and HBZ performed the experiments, analyzed data, and wrote the paper. All authors reviewed the manuscript and provided the input.

---

**Fig. S1.**
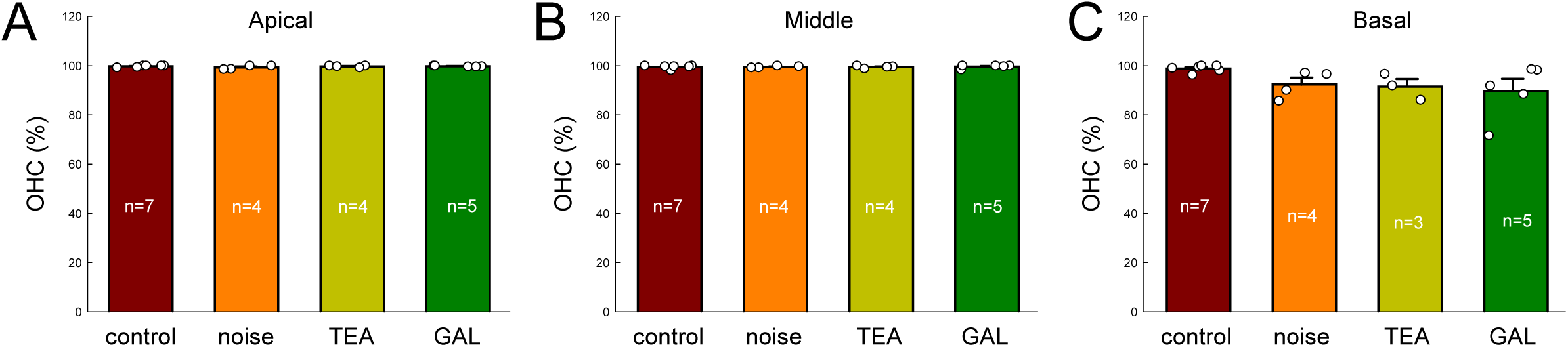
Hair cell degeneration in the experimental groups. Outer hair cells (OHCs) were accounted at the apical, middle, and basal turn in the control, noise, post-TEAx3d, and GALx3d groups. There are no significant OHC degenerations among these experimental groups.

